# A learned map for places and concepts in the human MTL

**DOI:** 10.1101/2020.06.15.152504

**Authors:** Nora A. Herweg, Lukas Kunz, Daniel R. Schonhaut, Armin Brandt, Paul A. Wanda, Ashwini D. Sharan, Michael R. Sperling, Andreas Schulze-Bonhage, Michael J. Kahana

**Affiliations:** Computational Memory Lab, Department of Psychology, University of Pennsylvania, Philadelphia, PA, USA; Department of Neuropsychology, Institute of Cognitive Neuroscience, Faculty of Psychology, Ruhr University Bochum, Germany; Department of Biomedical Engineering, Columbia University, New York, NY, USA; Epilepsy Center, Medical Center – University of Freiburg, Faculty of Medicine, Freiburg, Germany; Department of Neurosurgery, Thomas Jefferson University Hospital, Philadelphia, PA, USA; Department of Neurology, Thomas Jefferson University, Philadelphia, PA, USA

**Author notes:** Correspondence: Dr. Nora A Herweg, or Dr. Michael J Kahana.

## Abstract

Distinct lines of research in both humans and animals point to a specific role of the hippocampus in both spatial and episodic memory function. The discovery of concept cells in the hippocampus and surrounding medial-temporal lobe (MTL) regions suggests that the MTL maps physical and semantic spaces with a similar neural architecture. Here, we studied the emergence of such maps using MTL micro-wire recordings from 20 patients navigating a virtual environment featuring salient landmarks with established semantic meaning. We present several key findings: The array of local field potentials in the MTL contains sufficient information to decode subjects’ instantaneous location in the environment. Closer examination revealed that the field potentials represent both the subjects’ locations in virtual space and in high-dimensional semantic space. We further show that both spatial and semantic representations strengthen over time. This learning effect appears as subjects increase their knowledge of the environment’s spatial and semantic layout. Similarly, we observe a learning effect on temporal sequence coding. Over time, field potentials come to represent future locations, even after controlling for spatial proximity. This predictive coding of future states, more so than the strength of spatial representations per se, explains variability in subjects’ navigation performance. Our results thus support the conceptualization of the MTL as a memory space, building semantic knowledge, spatial and non-spatial, from episodic experience to plan future actions and predict their outcomes.

**Significance statement:** Using rare micro-wire recordings, we studied the representation of spatial, semantic and temporal information in the human MTL. Our findings demonstrate that subjects acquire a cognitive map that simultaneously represents the spatial and semantic relations between landmarks. We further show that the same learned representation is used to predict future states, implicating MTL cell assemblies as the building blocks of prospective memory functions.

## Introduction

Ever since Tolman first introduced the idea of a cognitive map (Tolman, 1948), researchers have debated its scope beyond the domain of physical space (Bellmund et al., 2018). If place cells in the rodent hippocampus act as pointers in a spatial cognitive map (O’Keefe, 1976; O’Keefe and Nadel, 1978), how do they relate to the episodic and semantic memory deficits seen in human patients with lesions or resections of the hippocampus and surrounding medial temporal lobe (MTL) cortex (Scoville and Milner, 1957)? O’Keefe and Nadel (1978) discussed the possibility that the human MTL maps semantic space like it maps physical space, but it was not until decades later that researchers discovered hippocampal concept cells (Quiroga, 2012; Reber et al., 2019) which fire for particular concepts in high-dimensional semantic space. Finally, time cells fire at particular times in a sequences of events (MacDonald et al., 2011; Eichenbaum, 2014; Umbach et al., 2020; Reddy et al., 2021). Together, these cell provide all components for the formation and retrieval of episodic memories, by defining “what” happened “where” and “when” (Tulving, 2001; Miller et al., 2013; Kunz et al., 2021).

However, few studies have studied spatial, semantic and temporal representations in combination. Moreover, it remains largely unclear how these different representations change during learning and how they relate to behavior. Here, we address this gap by analyzing micro-wire recordings from the human MTL. Patients who were undergoing clinical seizure monitoring learned to navigate a virtual environment, while instructed to successively find different target stores. Using a multivariate decoding approach, we asked whether the MTL would independently represent spatial, temporal and semantic information while subjects moved through the virtual city. In addition, we aimed to characterize learning-related changes to these representations. Whereas place fields in rodents navigating small vista-spaces become established within few minutes (Frank et al., 2004), we reasoned that representational changes in a complex environment with several distinct start and target locations may occur on a slower timescale. Finally, we asked how representations of space and temporal sequences relate to subjects’ behavior. Navigation performance in a novel environment tends to be variable and improve over time (Manning et al., 2014). However, it remains unclear how variability in behavior relates to variability in neural activity in human MTL populations.

To address these questions, we first trained a location decoder on the spectral features of the local field potential (LFP). The micro LFP is particularly well suited for this approach. It can be recorded on a large number of MTL channels (i.e. 8 micro-wires extending from the tip of each MTL depth electrode, even in the absence of large isolated spikes), while also providing high temporal stability, in particular across recording sessions. We trained our decoder to predict the subject’s location, defined as the label of the target store closest to the subject’s instantaneous position. We chose to define location in this manner to capture both semantic and spatial aspects of the subject’s position: each landmark is associated with a particular location in space and also has a predefined semantic meaning. We can thus derive a measure of representational content from the probabilities that the classifier assigns to each store: if the MTL represents the subject’s spatial location, the probabilities assigned to each store should depend on the subject’s spatial distance to that store; if the MTL represents the subject’s location in semantic space, the probabilities assigned to each store should depend on the semantic distance between the subject’s current location (nearest store) and each respective store.

Using a similar rationale, we also assessed the representation of temporal information. If the MTL tracks the sequence of past events (Howard and Kahana, 2002; MacDonald et al., 2011; Hsieh et al., 2014), the probabilities assigned to each store should depend on the time elapsed since the subject passed that store. We assumed, that if subjects learned the associations between the stores, they would also predict upcoming locations along their path. If these predictions are likewise represented in the MTL, the probabilities assigned to each store should depend on the time that will pass before a subject reaches a store.

## Results

We analyzed MTL micro-wire recordings from 20 patients undergoing clinical seizure monitoring who navigated a virtual city to deliver objects to different target stores **(Figure 1a-c)**. Subjects completed an average of 11.6 delivery days (min: 2, max: 22), during each of which they navigated to a random series of 13 different stores. We quantified subjects’ navigation performance using a ratio between their path length for each delivery and the shortest available path for that delivery. Over time (i.e. delivery days), subjects became more efficient at navigating to their target stores (χ^2^_(1)_ = 7.77, p = 0.005), confirming that they acquired knowledge for the spatial relations between the target stores.

**Figure 1.**
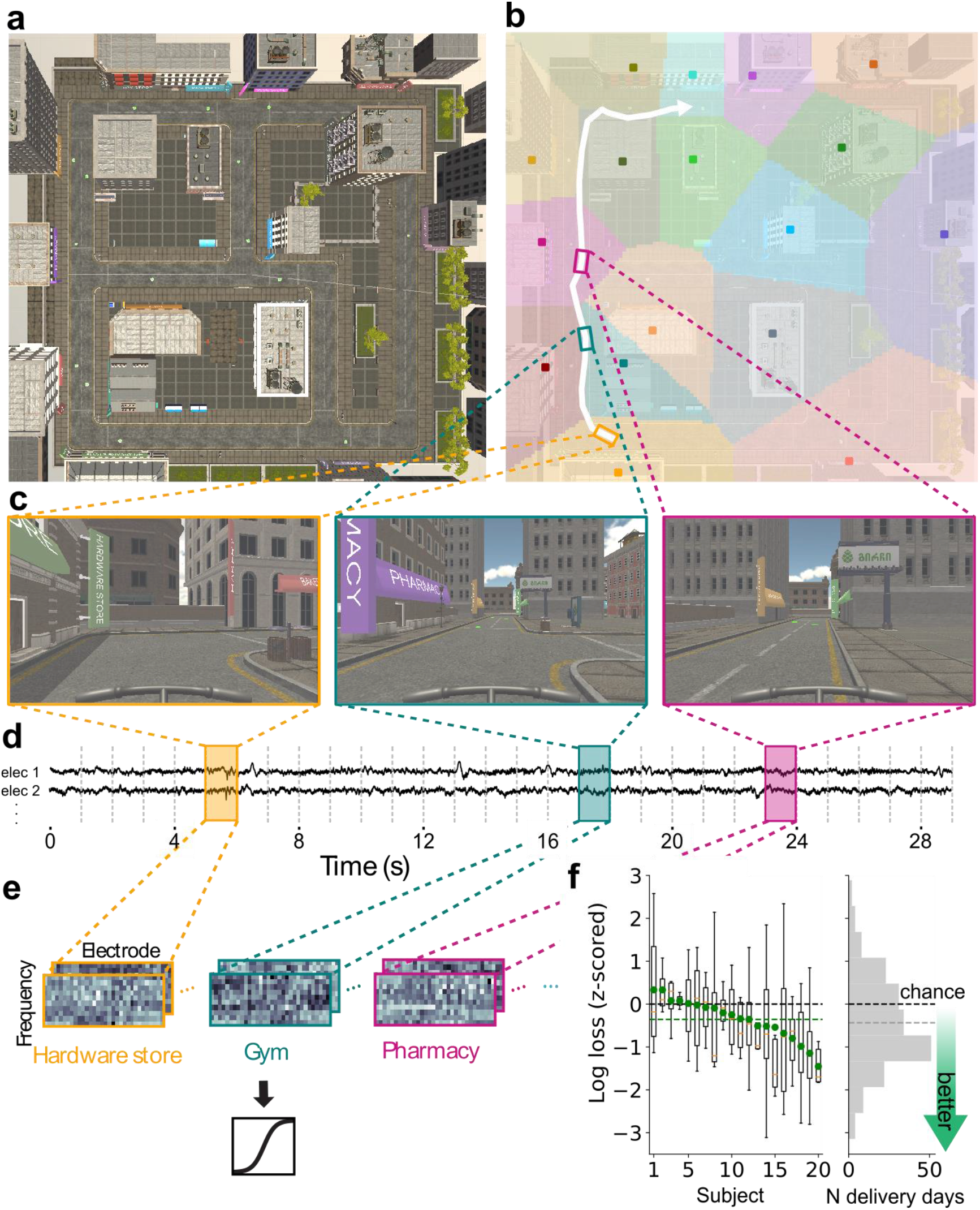
Multivariate decoding of location. a) Subjects navigated to a series of target stores in a virtual city. b) The target stores served as meaningful labels for different locations in the environment. Specifically, we assigned each coordinate in the environment to the store with the shortest Euclidean distance to that coordinate. The resulting borders are depicted along with a subject’s example path from the hardware store to the bike shop (white arrow). c) Snapshots taken during navigation for the three exemplary locations highlighted with colored rectangles, which were labelled as hardware store (yellow), gym (teal; note that the pharmacy is in view despite being further away) and pharmacy (pink). d) We extracted spectral power in 10 log-spaced frequencies between 3 and 200 Hz for each 1 second segment of continuous micro-wire LFP recordings. e) The N_frequency_ x N_electrode_ LFP power matrices were used as inputs to a multinomial logistic regression classifier predicting the subject’s instantaneous location for each 1 s epoch. f) We used logarithmic loss to assess decoding performance for each delivery day. A log loss of 0 indicates perfect classification performance with no uncertainty (i.e., a consistent classifier output of 100% for the true class). We z-scored classification performance for each delivery day relative to a chance distribution that was derived from classifiers trained on randomly shifted training labels. A z-score of 0 (black line) indicates chance performance, lower values indicate performance that is better than chance. The left panel shows the distribution of z-scored performance across delivery days (fliers not shown) for each subject, along with the subject-specific average (green points). Average z-scored performance across subjects (green line, left panel) was significantly better than chance. The right panel shows the aggregate distribution and average (grey line) across all delivery days.

We used a multinomial logistic regression classifier to decode location from LFP spectral power. To train our decoder, we segmented the LFP into 1s epochs and labeled each epoch with the target store that was closest to the subject’s instantaneous position (**Figure 1b-e**). We assessed significance of our model using a permutation procedure in which we trained “chance” classifiers on shuffled training labels (see Methods).

### LFP spectral power provides sufficient information to decode location

We find that the LFP contains sufficient information to predict the virtual location of human subjects with above chance accuracy (**Figure 1f**; t_(19)_ = −3.33, p = 0.003, Cohen’s d = −0.75; 10 log-spaced frequencies between 3 and 200 Hz). To determine whether decoding performance varied by MTL sub-region or frequency band, we trained separate classifiers on micro-wire bundles located in the hippocampus (N = 17 subjects), parahippocampal gyrus (N = 12 subjects) or amygdala (N = 12 subjects). Similarly, we trained separate classifiers on low-frequency (<30 Hz) or high frequency (>30Hz) features only. When compared to chance, decoding was significant in the hippocampus (t_(16)_ = −2.56, p = 0.02, Cohen’s d = −0.62; parahippocampal gyrus: t_(11)_ = - 2.07, p = 0.06, Cohen’s d = −0.60; amygdala: t_(11)_ = −1.13, p = 0.28, Cohen’s d = −0.33) and for high-frequency features (t_(19)_ = −3.83, p = 0.001, Cohen’s d = −0.86; low frequency: t_(19)_ = −0.65, p = 0.52, Cohen’s d = −0.15). However, when directly assessing the effects of brain region and frequency band, there was no effect of brain region (χ^2^_(2)_ = 2.58, p = 0.28; likelihood-ratio test comparing mixed-effects models, see Methods) and no effect of frequency band (t_(19)_ = 1.13, p = 0.27, Cohen’s d = 0.36) on classifier performance. We therefore focused all following analyses on the LFP-based decoding model trained on all frequencies and MTL sub-regions.

Having shown that we can predict the subject’s location from LFP spectral power, we next asked whether the MTL represents locations in physical or in semantic space. We define the former using the landmarks’ (i.e. the target stores’) locations in the virtual environment and the latter using their locations in word2vec space (Mikolov et al., 2013). Since all target stores had established semantic meaning, our model could achieve high performance based on either type of representation; the LFP could be representing either the subject’s spatial distance or their semantic distance to the stores (e.g. “at the landmark with coordinates (x_i_,y_i_), which is spatially proximate to the landmark with coordinates (x_j_,y_j_)” versus “at the pizzeria, which is semantically similar to the bakery”).

### Representation of virtual space strengthens over time

First, we examined representations of virtual space, using linear mixed effects models (see Methods). Specifically, we analyzed the distribution of classifier output probabilities for all stores as a function of the subject’s instantaneous spatial distance to those stores (Note that this analysis included only “incorrect” stores, thus excluding the one store closest to the subject’s instantaneous position). If the MTL represents spatial relations, the classifier should assign high probabilities to spatially proximate locations and low probabilities to spatially distant locations (**Figure 2a**; Note that this is independent of classifier performance, which is calculated based on the probabilities assigned to the correct store). We find that this is the case: Specifically, we observed a significant effect of the spatial distance between a given store and the subject’s true location on classifier output probabilities for that respective store (**Figure 2b**; z = −4.77, χ^2^_(1)_ = 22.73, p < 0.001).

**Figure 2.**
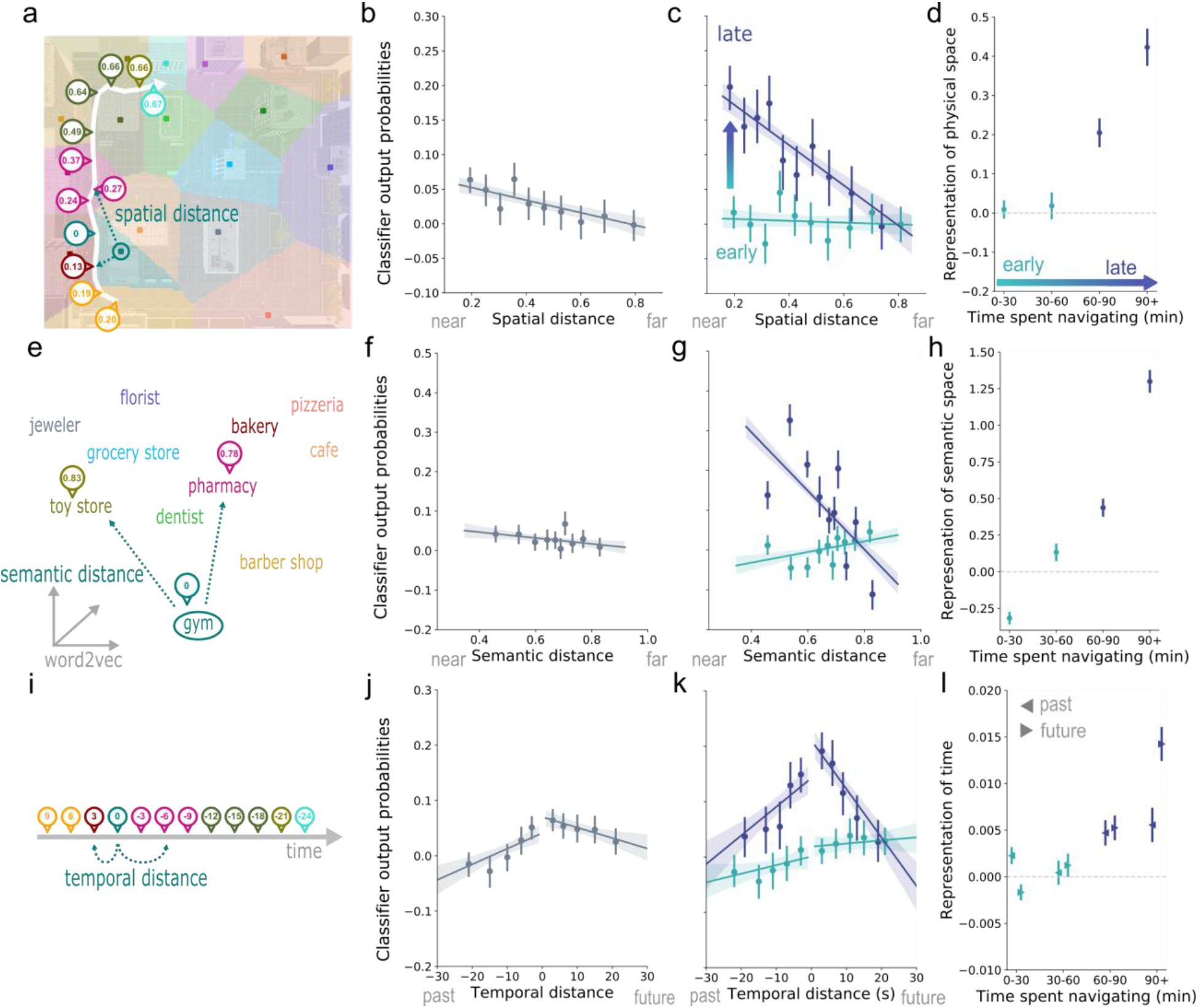
Representations of space, time and semantics strengthens with experience. a) A subject’s spatial distance from an example store, the gym, changes as the subject navigates. If the MTL represents the stores’ spatial locations, the classifier’s output probabilities for a given store should decrease with the subject’s distance to that store. b) We find that this is the case: classifier output probabilities decrease with the spatial distance between a given store and the subject’s true instantaneous position. Data are visualized after removing the estimated random effect for each subject. Error bars in panels b,c,f,g,j, and k depict a 95% confidence interval around the mean. c) The effect of spatial distance strengthens over time, suggesting that spatial knowledge manifests as increased representational similarity between nearby locations or an increased tendency for the MTL to represent proximate over distant spatial locations. For visualization, data was split into an “early” and a “late” phase of navigation at the 60 minute mark. d) Plotting the effect (i.e. slope) for four smaller time bins, reveals strengthened representations of space after 60 minutes that further increase after 90 minutes. Plotted are inverted slopes, which were estimated using linear effects models on the data from each bin. Positive values indicate a slope in the expected direction (i.e. higher classifier output for spatially proximate locations). Note how the average slope for early and late time periods matches the slope of the regression line depicted in panel c. Error bars in panels d, h, and l depict the standard error of the parameter estimate. e) Subjects move through semantic space, as they travel from store to store. In this example, the subject is semantically closer (in word2vec space) to the gym when located at the pharmacy compared with the toy store. We assessed the effect of semantic distance on classifier output for each store. f) Classifier output probabilities are high for stores that are semantically similar to the store at which the subject is currently located and low for stores that are semantically dissimilar to the subject’s current location. g,h) Again, the effect of semantic distance strengthens over time, suggesting that the MTL increasingly represents the subject’s location in semantic space. i) Finally, we asked how classifier output for a given stores relates to the time that has passed since the subject has visited that store (negative distance), or to the time that will pass until the subject arrives at that store (positive distance). j) Classifier output is generally higher for stores that a subject is about to visit than for those that have been visited in the recent past. k,l) Like for spatial and semantic distance, the effect of temporal distance increases over time. This increase is stronger for stores on a subject’s future path, meaning that with experience the MTL more strongly predicts a subject’s future trajectory.

**Figure 3.**
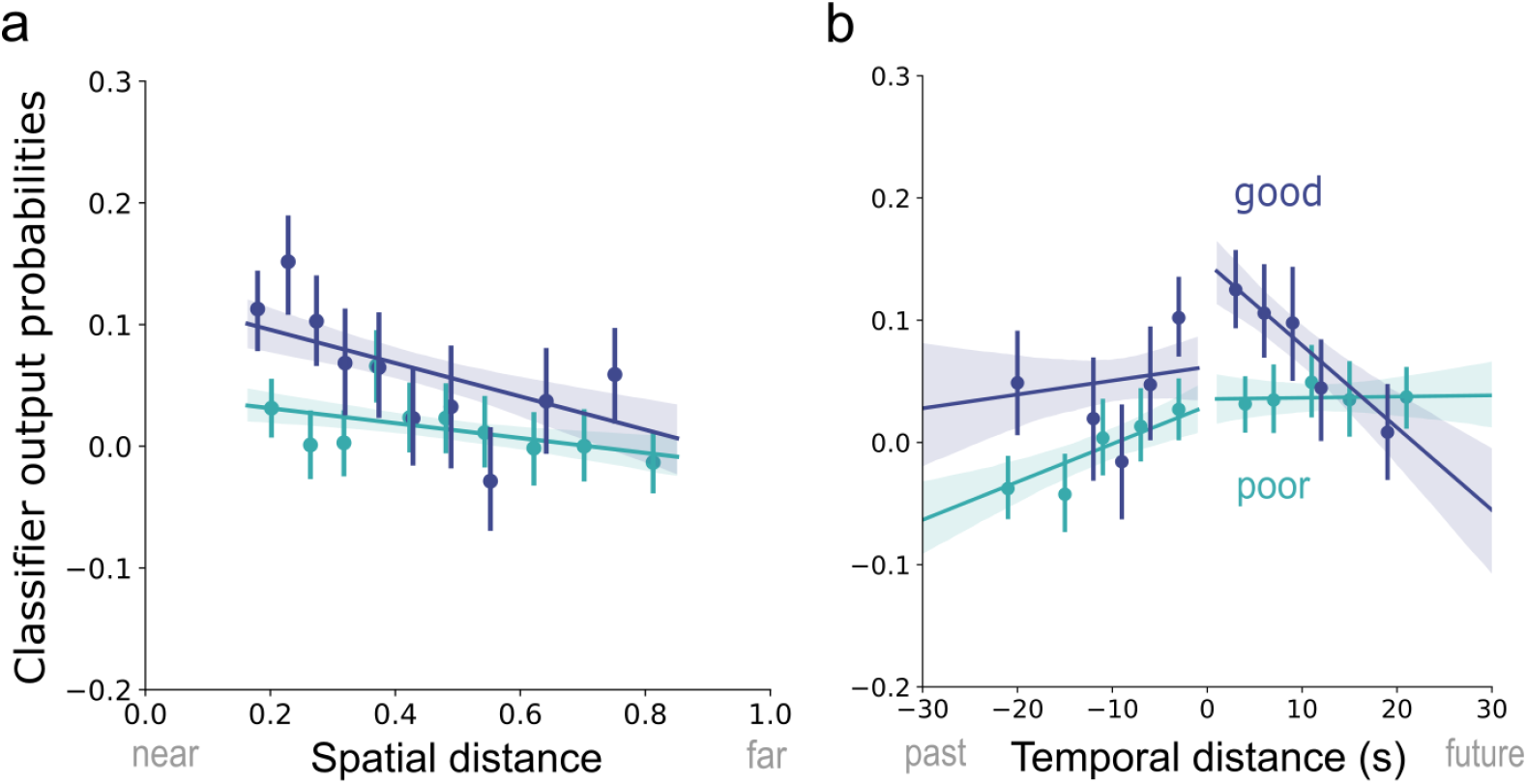
Representation of time linked to navigation performance. a) Representations of space (i.e. the effect of spatial distance on classifier output) may not only change over time but may also be linked to navigation performance (visualized here for “good” vs. “poor” performance trials). We find that this is the case when assessing spatial distance individually but not when controlling for temporal distance. Data are visualized after removing the estimated random effect for each subject. Error bars depict a 95% confidence interval around the mean. b) The interaction between navigation performance and temporal distance, on the other hand, remains significant even after accounting for spatial distance and is particularly strong for upcoming locations. Note how the slope is steeper for “good” compared with “poor” performance trials, in particular on the right-hand side, displaying future locations. While general knowledge of the city’s spatial layout is a prerequisite for spatial planning, it seems that the success of this process on individual trials, leading to robust prediction of upcoming locations, most closely relates to subjects’ navigation performance.

In our task, spatial and temporal distance were correlated (r_spatial,temporal_ = 0.71), but semantic distance was uncorrelated with both of them (_rsemantic,spatial_ = 0.05, r_semantic,temporal_ = 0.03). To control for the positive correlation between spatial and temporal distance, we ran a second model, in addition to the “individual model” reported above, that included both spatial and temporal distance as a predictor. While this approach slightly complicated the interpretation of null effects (they may be true null-effects or observed due to the shared variance in space and time), we reasoned that it provides a conservative means to assess the independent contributions of each predictor: If spatial distance explains variability in classifier output independent of time, then it should remain significant in the “joint model”. Indeed, the effect of spatial distance was significant in both models (joint model: z = −2.89, χ^2^_(1)_ = 8.37, p = 0.003). Taken together, these results indicate that the LFP represents the subject’s spatial location within a broad, spatial map of the environment. Moreover, this effect remained significant even after accounting for correlations between movement through time and space.

Next, we examined whether the neural representation of spatial relations exhibits a learning effect. Having shown that subjects navigate more efficiently as they acquire spatial knowledge of the environment, we expected this behavioral improvement to be reflected in a change of the representational structure in the MTL. We find that the effect of spatial distance on classifier output probabilities strengthens over time, as shown by a significant interaction of spatial distance and epoch number (individual: z = −6.71, χ^2^_(1)_ = 45.0, p < 0.001; joint: z = −3.58, χ^2^_(1)_ = 12.85, p < 0.001). This effect is illustrated in **Figures 2c**, which shows classifier output probabilities as a function of spatial distance for an “early” and a “late” phase of navigation, split at the 60 minute mark. **Figure 2d** displays the slope of this regression for four smaller time bins (30 minute increments), revealing an initial increase after 60 minutes, that further strengthens after 90 minutes. This finding provides evidence that spatial representations in the human MTL change over the course of learning: Specifically, spatial knowledge manifests as increased representational similarity between nearby locations or an increased tendency for the MTL to represent proximate over distant spatial locations.

### Representation of semantic space strengthens over time

We next asked whether MTL activity reflects the semantic structure of the environment. To this end, we calculated the distances in word2vec (Mikolov et al., 2013) semantic space between all target stores and assessed their relationship with the classifier’s predictions (**Figure 2d**). We find that the classifier assigns higher probabilities to stores that are semantically related to the store at which the subject is currently located (e.g. the pizzeria while the subject is located at the bakery; **Figure 2e**; z = −2.03, χ^2^_(1)_ = 4.13, p = 0.04). The MTL, thus, not only represents the subject’s location in virtual space but also the subject’s location in semantic space. This finding complements recent reports of concept coding in the human MTL (Quiroga, 2012; Constantinescu et al., 2016).

Whereas we had a strong expectation that subjects’ spatial knowledge would increase over time, one may argue that semantic information (e.g. the concept of a bakery) should have been firmly established through prior experience, and would thus not be expected to change over the course of the experiment. However, it is possible that MTL neural activity adapts to the specific semantic subspace spanned by our task, reflecting semantic similarity on fewer relevant dimensions. To evaluate these alternative predictions, we asked whether the degree to which the MTL represents the subject’s semantic distance to each store changes over time (i.e. the distance between the nearest store and all other stores at each point in time). As with spatial information, we find that the link between semantic distance and classifier output strengthens over time (**Figure 2f**; z = - 16.16, χ^2^_(1)_ = 260.87, p < 0 .001). This means that the neural similarity between related task-relevant concepts (such as the pizzeria and the bakery) increases throughout the experiment, more so than the neural similarity between task-relevant unrelated concepts.

### Representation of temporal sequence strengthens over time

Finally, we asked whether the LFP represents the temporal sequence of visited locations, when controlling for the spatial distance between them. We expected that representations of stores would linger after they were visited and that the MTL would come to predict upcoming locations along a subject’s future trajectory. To address this question we assessed the link between classifier output for each store and the temporal distance between the current time point t_0_ and the last time a subject had visited that store t_0-j_ (past) or the next time a subject was going to visit that store t_0+i_ (future; **Figure 2g**). We find that temporal distance does affect classifier output, when assessed individually (**Figure 2h**; z = −3. 91, χ^2^_(1)_ = 15.29, p < 0 .001), but not when controlling for spatial distance (z = −0.48, χ^2^_(1)_ = 0.23, p = 0.63). However, we also find that the effect of time is asymmetric, in that future visits are more strongly represented than those in a subject’s past (individual: z = 5.57, χ^2^_(1)_ = 30.98, p < 0.001; joint: z = 5.61, χ^2^_(1)_ = 31.45, p < 0.001). So even though there seems to be no fine grained sequence information, the MTL broadly differentiates between upcoming and recently visited locations.

Even though the MTL does not represent fine grained sequence information when including all data, we asked whether sequence coding, and in particular the prediction of upcoming locations, gets stronger over time. Because subjects need to know the city’s spatial layout before being able to plan routes and predict future states, we hypothesized that temporal sequence coding may emerge later in the experiment. We could confirm this prediction, in particular, for locations on a subject’s future trajectory, as indicated by a three-way interaction of absolute temporal distance, epoch number and future vs. past visits (**Figure 2i**; individual: z = −3.62, χ^2^_(1)_ = 13.11, p < 0.001). This effect holds when controlling for spatial distance (joint: z = −4.15, χ^2^_(1)_ = 17.22, p < 0.001). The MTL thus predicts upcoming locations more strongly, as subjects spend more time navigating the environment.

### Temporal sequence coding explains navigation performance

Having shown that representations of time and space get sharper over time, we wondered whether variance in the sharpness of these representations may explain subjects’ navigation performance. Do subjects navigate more efficiently, when MTL activity reflects the spatial layout of the environment and the particular temporal sequence of visited locations? We find that this is the case. While controlling for time on task, we observe a significant interaction between spatial distance and excess path ratio on classifier output probabilities (z = 2.57, χ^2^_(1)_ = 6.60, p = 0.01), suggesting that the MTL represents the city’s spatial layout more strongly at times when subjects navigate to their goal efficiently. Similarly, we find a significant interaction between temporal distance and excess path ratio on classifier output (z = 4.72, χ^2^_(1)_ = 22.26, p < 0.001), suggesting that temporal sequence is represented more strongly during deliveries with high navigation performance. Again, we repeated these analyses, including the other factor respectively, to see if temporal and spatial distances explain independent variance. When controlling for temporal distance, the effect of spatial distance disappears (z = −0.99, χ^2^_(1)_ = 0.99, p = 0.32). The effect of temporal distance, however, withstands correction for spatial distance (z = 4.09, χ^2^_(1)_ =16.73, p < 0.001), and is marginally stronger for locations that lie in the future compared to the past (three-way interaction, z = 1.61, χ^2^_(1)_ = 3.44, p = 0.06). Together these results suggest that the MTL function that is most strongly linked to subjects’ navigation performance is the simulation of specific future paths.

## Discussion

Here, we used a decoding-based approach to study neural representations of places, concepts and temporal sequences. As a first step, we trained a location decoder on the micro LFP while subjects navigated a virtual city. We find that the spectral features of the micro LFP provide sufficient information to decode subjects’ instantaneous location in the virtual city with above-chance accuracy. As such, our results provide the first demonstration of location decoding from local field potentials in the human MTL and they are in line with a prior study in rodents showing that hippocampal theta oscillations can predict rat position (Agarwal et al., 2014). Because research suggests that the human hippocampus does not exhibit theta oscillations in the same continuous way as that of rodents (Watrous et al., 2013; Aghajan et al., 2017) and since the number of micro-electrodes is limited by clinical considerations, we defined our feature set more broadly including multiple MTL subregions (hippocampus, parahippocampal gyrus, and amygdala) and frequency bands. We find that, when working with subsets of the data, classification did not differ between MTL regions or frequency bands. Similar findings have been obtained with human single unit recordings, showing that place-selective neurons are not confined to the hippocampus but can also be observed in the PHG and amygdala (Jacobs et al., 2013; Miller et al., 2013). It is, however, possible that the regional non-specificity can partly be explained by the difficulty of precisely localizing micro-wire electrodes in human subjects.

We used our trained decoding model to characterize the representations that underlie successful decoding. As subjects move around the virtual environment, navigating from store to store, they change their virtual location, but, since all stores have pre-established semantic meaning, they also change their semantic location (e.g. activating the concept “bakery”). Our decoding model has been trained in a way that is blind to this distinction (i.e. using the stores as labels for different parts of the environment), meaning that it could achieve above-chance performance based on either type of representation, spatial or semantic. By analyzing the classifier’s output probabilities to each store at every time point, we found that the MTL represents locations in both virtual space (similar neural representations for spatially proximate stores) and semantic space (similar neural representations for semantically similar stores).

Considering virtual space, we show that the classifier’s output probabilities depend on the spatial distance between a subject’s true location and each store. The classifier, on average, assigns a high probability to the store that is closest to the subject’s true location. For other locations, the classifier assigns decreasing probabilities to stores that are farther from the subject’s true location. The representational space spanned by MTL spectral features, hence, mirrors the virtual space spanned by the landmarks in the virtual city. These results resemble similar findings obtained with representational similarity analysis on fMRI data from the human hippocampus (Deuker et al., 2016). Moreover, we observed evidence that the spatial coding effect strengthens over time, providing a window into the learning related changes of representational structure in the MTL. The more time subjects have spent navigating the virtual city, the more strongly the MTL represents the city’s spatial layout. Whereas place fields in rats foraging in small open environments are known to stabilize within a few minutes of entering the environment (Wilson and McNaughton, 1993; Frank et al., 2004), the changes we observed here occurred over a duration of more than an hour. This time scale is plausible given the complexity of environment and the fact that we observed concurrent increases in subjects’ spatial navigation performance.

We find that in addition to representations of virtual space, the MTL also tracks subjects’ movement through semantic space. Subjects’ virtual movements from store to store can be thought of as movement through high-dimensional semantic space, activating the concept of each store as they pass it. Evidence for this interpretation comes from the fact that the classifier output probabilities for each store are related to that store’s semantic distance from the subject’s true location. For instance, while the subject is near the bakery, the classifier tends to assign high probabilities to related stores such as the pizzeria. And even though the concepts used in this study were familiar to subjects at the outset (e.g. subjects did not have to newly learn the concept “bakery”), we observed that the link between classifier output and semantic distance increased over time, just like it did for spatial distance. Our finding suggests that concept coding in the MTL is dynamic. Specifically, we interpret this effect as a “zooming in” in high-dimensional semantic space. As the subject acquires knowledge of the environment’s specific semantic subspace, MTL activity seems to reflect the semantic relations in this space on a reduced number of task-relevant dimensions.

Finally, we find that the MTL represents the temporal sequence of visited locations, more so at the end then the beginning of the experiment. This effect is particularly strong in the forward direction, meaning that the MTL increasingly predicts locations that a subject is about to pass in the near future. The more knowledge subjects acquire about the virtual city’s spatial layout, the more they are capable of planning specific trajectories when looking for their target store, including the specific sequence of stores that they will pass on their way. And intriguingly, it is this predictive signal that explains subjects’ navigation performance for individual deliveries, more so than it is a general representation of the city’s spatial layout. This finding implicates the MTL not only in the formation and storage of a cognitive map, but also in accessing the map to predict future states.

## Conclusions

Our findings demonstrate that field potentials in the human MTL jointly represent a subject’s virtual spatial location, the semantics associated with that location, and a subject’s temporal trajectory. The representation of space, semantics, and time becomes more pronounced with experience, indicating that the MTL stores a learned map of virtual and local semantic space that can be used to predict future trajectories.

## Materials and Methods

### Participants

20 patients with medication-resistant epilepsy undergoing clinical seizure monitoring at Thomas Jefferson University Hospital (Philadelphia, PA, USA) and the University Clinic Freiburg (Freiburg, GER) participated in the study. The study protocol was approved by the Institutional Review Board at each hospital and subjects gave written informed consent. Subjects were implanted with Behnke-Fried macro-micro depth electrodes (AdTech, Racine, WI) in the MTL. The location of these electrodes was determined based on clinical considerations.

### Experimental design and task

Subjects played the role of a bicycle courier in a hybrid spatial-episodic memory task, delivering parcels to stores located within a virtual town (consisting of roads, stores, and task-irrelevant buildings; **Figure 1**). Subjects completed a variable number of experimental sessions (min: 1, max: 4, mean = 2.45). Each experimental session consisted of a series of delivery days. On each delivery day, subjects were first instructed to deliver a series of objects to specific stores in the city (see below for details). Then, after navigating to the last store, subjects were tested on their memory for these objects. In the current study, we used data from the navigation phases only; data from the recall periods for an overlapping sample of subjects has been reported elsewhere (Miller et al., 2013; Herweg et al., 2020). Subjects completed slightly different versions of this paradigm, the details of which are described in the following paragraphs. We believe these task adaptations to be negligible in terms of the scope of the paper and report all results collapsed over all subjects/versions. The tasks were programmed and displayed to subjects using the Panda Experiment Programming Library (Solway et al., 2013), which is a Python based wrapper around the open source game engine Panda3d (with 3D models created using Autodesk Maya™) or the Unity Game Engine (Unity Technologies, San Francisco, CA).

Prior to starting the first delivery day, subjects viewed a static or rotating rendering of each store in front of a black background (later referred to as ‘store familiarization’). Each store had a unique storefront and a sign that distinguished it from task-irrelevant buildings. Each delivery day consisted of a navigation phase (**Figure 1**) and a recall phase (for which no data is reported here). For the navigation phase, 13 stores were chosen at random out of the total number of 16 or 17 stores. Subjects were informed about their upcoming goal by on-screen instructions (e.g. “Please find the hardware store.”) and navigated to each store using the joystick or buttons on a game pad. Location-store mappings in the town were fixed for 4 subjects and random for 16 subjects (the layout was always fixed across experimental sessions; i.e. each subject experienced the same town layout across sessions). Upon arrival at the first 12 stores, subjects were presented with an audio of a voice naming the object or an image of the object they just delivered. Object presentation was followed by the next on-screen navigation instruction. Upon arrival at the final store, where no item was presented, the screen went black and subjects heard a beep tone. After the beep, they had 30 or 90 s to recall as many objects as they could remember in any order. A final free recall phase followed on the last delivery day within each session.

To ensure that subjects did not spend extensive time searching for a store, waypoints helped a subset of subjects (N = 4 subjects) navigate: Considering each intersection as a decision point, arrows pointing in the direction of the target store appeared on the street after three bad decisions (i.e. decisions that increased the distance to the target store). In a different version of the task (N = 5 subjects), subjects had to complete a pointing task prior to navigation to each store. From their current location, they were asked to use the game pad to point an arrow in a straight-line path to where they thought the target store was located. This task served a similar purpose as the waypoints, since it provided subjects with feedback on the correct direction of the target store prior to navigation.

Most subjects (N = 16) completed an initial learning session prior to the first delivery day session, in which the store familiarization phase was followed by a ‘town familiarization’ phase. Here, subjects were instructed to navigate from store to store without delivering parcels, or later recalling objects, visiting each store three times in pseudo-random order (each store was visited once, before any repeat visit). LFP data during the learning session was available only for a subset of subjects (N = 4). For these subjects, data from the very first round of navigation was not used in the analyses. For some subjects (N = 4), a single town familiarization trial was repeated at the beginning of all following sessions prior to the first delivery day.

### Behavioral analyses of navigation performance

Behavioral data were analyzed using Python version 3.6. As a measure of navigation performance, we computed the excess path length ratio. This is the actual length of a subject’s path (i.e. from the initial instruction “please find the hardware store” to arrival at that store) divided by the shortest possible path between the start and arrival locations. A value of 1 indicates perfect performance, higher values indicate a suboptimal path. We used a mixed linear model with a random subject intercept to assess the effect of delivery day number on navigation performance. We performed a likelihood ratio test between a model including the fixed effect of delivery day number and an intercept-only model to test for a significant main effect.

### Micro-wire data acquisition and preprocessing

LFPs were recorded from the inner micro-wire bundle of Behnke-Fried macro-micro depth electrodes (AdTech, Racine, WI) located in the MTL (including hippocampus, N = 17 subjects; parahippocampal gyrus, N = 12 subjects and amygdala, N = 12 subjects). Data were recorded at a sampling rate of 20,000 or 30,000 Hz using the NeuroPort™ system (Blackrock, Salt Lake City, UT), a Neuralynx system (Neuralynx, Bozemann, MT) or an inomed system (inomed, Emmendingen, Germany). Data from the eight recording wires on each bundle were referenced online to a ninth designated reference wire. In cases where the reference wire was suspected to have low signal quality, one of the regular recording wires was used as an online reference instead. No re-referencing was performed offline. Coordinates of the radiodense macro-electrode contacts were derived from a post-implant CT or MRI scan and then registered with the pre-implant MRI scan in MNI space using SPM or Advanced Normalization Tools (ANTS)(Avants et al., 2008). For 7 subjects, micro-wire bundles were localized manually by a neurologist or a radiologist, for the other 13 subjects, micro-wire bundles were localized by extrapolating 0.5 cm (the approximate length the micro-wires extend from the tip of the electrode) from the location of the most distal macro-electrode contact. Labels were derived using the probabilistic Harvard Oxford atlas with a threshold of 25%.

LFP data were analyzed using Python version 3.6 along with the Python Time Series Analyses (ptsa; https://github.com/pennmem/ptsa_new) and MNE (Gramfort et al., 2014) software packages. LFP data were aligned with behavioral data via pulses sent from the behavioral testing laptop to the recording system. A zero-phase notch filter was applied to filter line noise at 50 Hz (for data recorded in Germany) or 60 Hz (for data recorded in the US) and harmonics up to 240 Hz (stopband width: frequency/200). To extract LFP power, continuous data were down-sampled to 1000 Hz and convolved with complex Morlet wavelets (5 cycles) for 10 approximately log-spaced frequencies between 3 and 200 Hz. Some frequencies were slightly shifted to avoid overlap with line noise frequencies (3, 5, 8, 12, 19, 31, 40, 79, 130, 210 Hz). After convolution, data were log-transformed and averaged over 1 s non-overlapping epochs.

### Multivariate classification

We used L2 penalized multinomial logistic regression to decode a subject’s current location across these 1 s navigation epochs. As features we used the N_electrode_ * N_frequency_ LFP power feature matrices. All epochs were labeled using the stores in the environment (N classes = N stores). Specifically, for each of the 1 s epochs we determined the subject’s current location by finding the store with the shortest Euclidean distance to the subject’s instantaneous position. Since these labels may change over successive samples within a 1 s epoch, we assigned each epoch the most frequent label out of the set of labels for that epoch.

We assessed classifier performance using a nested leave-one-delivery-day-out cross-validation procedure (each delivery day includes navigation to 13 stores and lasted several minutes). To make sure that test and training data were sufficiently separated in time, we never tested the classifier on data from the initial learning session. Because navigation in the learning session was continuous and not separated from one another by a recall phase or a short break, these data were only used for classifier training. Nested cross validation was used to optimize the penalty parameter C using a grid search with 20 log spaced values between log_10_(10^-4^) and log_10_(200). To this end, the training data within each outer cross-validation fold was again divided into inner leave-one-delivery-day-out cross-validation folds. The classifier was trained on each inner training set for all 20 C values. The optimal C for a given outer fold was chosen as the C that maximized classifier performance across the inner test sets. For each inner and outer fold, all data were z-scored with respect to the mean and standard deviation of the respective training set. Samples were weighted inversely proportional to their class frequency to avoid bias towards more frequently occurring classes.

We evaluated performance using logarithmic (or cross-entropy) loss (again weighting every sample inversely by its class frequency), which is defined as the negative logarithm of the probability assigned to the true class. Log loss therefore produces high values, when the assigned probability for the true class approaches zero (i.e. a confident incorrect classification) and values close to zero when the assigned probability for the true class approaches one (i.e. a confident correct classification). The value of log loss that corresponds to chance depends on the number of classes, i.e. stores. The number of stores in the environment was variable across subjects (N_stores_ = 16 or 17). We therefore used a permutation procedure (N_permutations_ = 200) to z-score the log loss for each delivery day against its individual chance distribution. Specifically, we repeatedly trained the classifier on a shuffled version of the training data (again using the nested cross-validation scheme outlined above). On each permutation, the true y-vector (i.e. location labels) was flipped and circularly shifted by a random number of elements in the vector. This procedure removes any true relation between features and to-be-predicted labels, while keeping the autocorrelation of the to-be-predicted labels unchanged. We then z-scored true classifier performance for each delivery day using the mean and standard deviation of that delivery day’s random distribution. z-scores < 0 indicate above-chance performance and z-scores > 0 indicate below chance performance.

A one-sample t-test across subjects was used to compare average z-scored classifier performance to chance. To evaluate the importance of activity from different MTL sub-regions, we additionally trained the classifier on a reduced feature set, which included only wire bundles in the hippocampus, parahippocampal gyrus or amygdala. Likewise, we trained separate classifiers on LFP power in frequencies below or above 30 Hz. We used a t-test across subjects to assess the effect of frequency band and a mixed linear model with a random subject intercept to assess the effects of sub-region on classifier performance. A mixed effects model is better suited for the latter effect because not all subjects had electrodes in all brain regions of interest. In the mixed model, we assessed significance of the single fixed effect (brain region) using a likelihood ratio test between the full model (main effect included) and an intercept-only model.

### Analyses of classifier output probabilities

Classifier output probabilities from the test sets were z-scored with respect to their permutation distribution (see above). We then used a mixed linear model with a random intercept for class (i.e., unique locations) nested in subject to assess the effect of spatial distance, temporal distance, semantic distance, as well as their interactions with epoch number or navigation performance on classifier output probabilities. We excluded from this analysis all probabilities for the subject’s true location, meaning that no statistical effect could be driven by a change in classifier accuracy. For each class (i.e. store) and each 1 s epoch, spatial distance was calculated as the Euclidean distance between the subject’s current position and the location of the respective store. Temporal distance was calculated as the absolute temporal distance in seconds between the current epoch and the last or next epoch during which the subject was or will be located at the respective store. An additional binary regressor was included to model past vs. future time points. Because there was less data for long temporal distances, we only included data points with an absolute temporal distance < 30 s. Semantic distance was calculated as the Euclidean distance in word2vec space (Mikolov et al., 2013) between the name of a given store and the name of the store at which the subject is currently located. We assessed significance of fixed effects using likelihood ratio tests between a full model (all main effects or all main effects and all interaction terms) and a reduced model (main effect or interaction effect in question removed).

## Data availability

Data that can be shared without compromising research participant privacy/consent will be available at http://memory.psych.upenn.edu/Electrophysiological_Data.

## Acknowledgements

This work was supported by German Research Foundation (DFG) grants HE 8302/1-1 to NAH and KU 4060/1-1 to LK, National Science Foundation (NSF) grant BCS-1724243 to MJK, National Institutes of Health (NIH) grants MH061975 and NS113198 to MJK and Federal Ministry of Education and Research (BMBF) grant 01GQ1705A to ASB. We thank Alison Xu, Zeinab Helili, Katherine Hurley, Deb Levy, Logan O’Sullivan, Ada Aka and Allison Kadel for help with data acquisition and post-processing, Jonathan Miller and Ansh Johri for their contributions to the task design, Corey Novich and Ansh Patel for programming the Unity based experiment, as well as Joel Stein, Rick Gorniak and Sandy Das for electrode localization support. We are most grateful to all patients and their families who selflessly volunteered their time to participate in this research.

## Author contributions

NAH and MJK designed research, NAH analyzed data, NAH and MJK wrote the paper, AB, LK, PAW and NAH collected data, AS-B, MRS, ADS recruited participants and performed clinical duties associated with data collection including neurosurgical procedures or patient monitoring, all authors reviewed and edited the manuscript.

